# MLL3/MLL4 Histone Methyltranferase Activity Dependent Chromatin Organization at Enhancers during Embryonic Stem Cell Differentiation

**DOI:** 10.1101/2021.03.17.435905

**Authors:** Naoki Kubo, Rong Hu, Zhen Ye, Bing Ren

## Abstract

MLL3 (KMT2C) and MLL4 (KMT2D), the major mono-methyltransferases of histone H3 lysine 4 (H3K4), are required for cellular differentiation and embryonic development in mammals. We previously observed that MLL3/4 promote long-range chromatin interactions at enhancers, however, it is still unclear how their catalytic activities contribute to enhancer-dependent gene activation in mammalian cell differentiation. To address this question, we mapped histone modifications, long-range chromatin contacts as well as gene expression in MLL3/4 catalytically deficient mouse embryonic stem (ES) cells undergoing differentiation toward neural precursor cells. We showed that MLL3/4 activities are responsible for deposition of H3K4me1 modification and formation of long-range enhancer-promoter contacts at a majority of putative enhancers gained during cell differentiation, but are dispensable for most candidate enhancers found in undifferentiated ES cells that persist through differentiation. While transcriptional induction at most genes is unaltered in the MLL3/4 catalytically deficient cells, genes making more contacts with MLL3/4-dependent putative enhancers are disproportionately affected. These results support that MLL3/4 contributes to cellular differentiation through histone-methyltransferase-activity dependent induction of enhancer-promoter contacts and transcriptional activation at a subset of lineage-specific genes.

## INTRODUCTION

Spatiotemporal gene expression in mammals is governed primarily by transcriptional enhancers, where binding of sequence-specific transcription factors (TFs) drives local chromatin changes by recruiting chromatin remodeling complexes such as SWI/SNF proteins and chromatin modifiers (Clapier and Cairns, 2009; Euskirchen et al., 2012; Heintzman et al., 2009; Long et al., 2016). Some of the most pronounced histone modifications found at enhancers include mono-methylation of histone H3 lysine 4 (H3K4me1) and acetylation of histone H3 lysine 27 (H3K27ac), which have been broadly utilized to identify and annotate enhancers in the genome (Andersson et al., 2014; Calo and Wysocka, 2013; Consortium, 2012; Creyghton et al., 2010; Rada-Iglesias et al., 2011; Shen et al., 2012; Shlyueva et al., 2014). Histone H3 lysine 4 mono-methylation at enhancers is catalyzed by the histone methyltransferases MLL3 and MLL4 (MLL3/4) (Herz et al., 2012; Hu et al., 2013; Lee et al., 2013; Wang et al., 2016), while H3K27ac is catalyzed by CBP/p300, recruitment of which could be facilitated by MLL3/4 (Jin et al., 2011; Lai et al., 2017). MLL3/4 play crucial roles in mammalian development. *Mll4* knockout in mice leads to embryonic lethality (Ashokkumar et al., 2020; Lee et al., 2013), and development of heart, adipose, muscle, and immune cells is severely impeded after *Mll3/4* depletion (Ang et al., 2016; Lee et al., 2013; Placek et al., 2017). Furthermore, mutations in *MLL3/4* genes are frequently observed in human cancers and developmental disorders (Ng et al., 2010; Parsons et al., 2011; Pasqualucci et al., 2011; Sze and Shilatifard, 2016; Will and Steidl, 2014). However, the role of MLL3/4 catalytic activity and MLL3/4-dependent H3K4me1 at enhancers is still incompletely defined.

A recent study demonstrated that catalytic inactivation of MLL3/4 causes loss of H3K4me1 at enhancers along with partial reduction of H3K27ac in mouse embryonic stem cells (ESCs), but with surprisingly minor effects on gene expression (Cao et al., 2018; Dorighi et al., 2017). Additionally, catalytic inactivation of Trr, the *Drosophila* homolog of MLL3/4, does not impede *Drosophila* development (Rickels et al., 2017). These observations raise the questions about the role of MLL3/4-depednent H3K4me1 in enhancer-dependent gene activation in general. We previously showed that MLL3/4 regulate chromatin organization at enhancers and modulate enhancer-promoter (E-P) contacts at the *Sox2* gene in mouse embryonic stem cells (Yan et al., 2018). However, the scope of MLL3/4-dependent histone methylation and its impact on E-P contacts and transcriptional programs of cellular differentiation required further investigation. To gain a better understanding of MLL3/4’s role at enhancers, it is essential to precisely determine how MLL3/4-dependent H3K4me1 regulates the dynamics of chromatin contacts between enhancers and promoters and expression of target genes (Deng et al., 2012; Gorkin et al., 2014). Here we used mouse ESCs with catalytically deficient MLL3/4 (hereafter referred to as dCD) (Dorighi et al., 2017; Li et al., 2016; Zhang et al., 2015) to delineate the role of MLL3/4 activities in histone H3K4 methylation at enhancers, E-P contacts, and transcriptional induction during ESC differentiation (Figure 1A, Table S1). As a model for cellular differentiation, we focused on retinoic acid (RA)-induced neural differentiation toward neural precursor cells (NPCs) (Methods) (Bain et al., 1995; Strubing et al., 1995).

**Figure 1.**
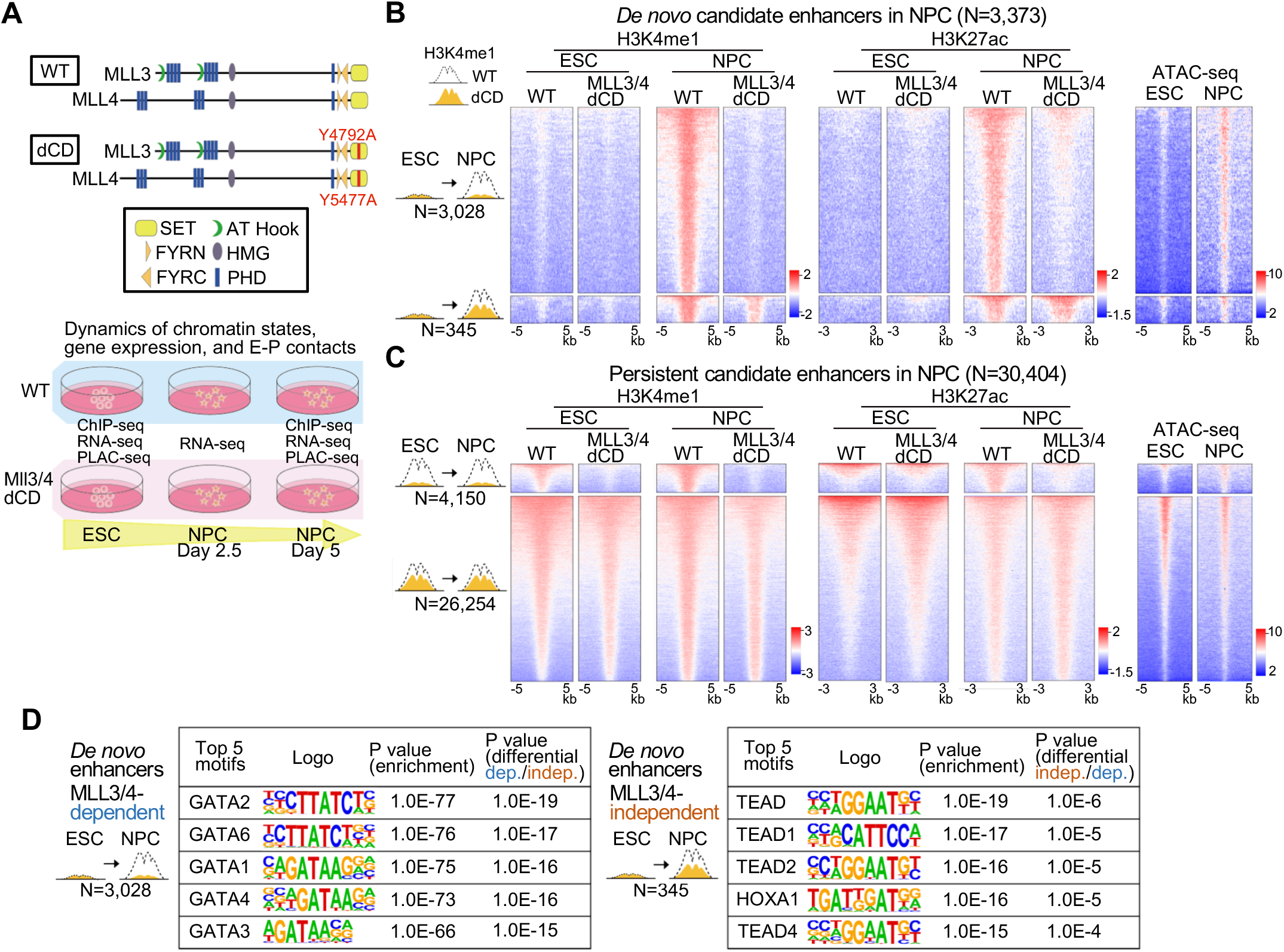
Loss of MLL3/4 catalytic activity leads to failure of accumulation of H3K4me1 at *de novo* enhancers during NPC differentiation of ESC. **(A)** Schematic representation of wild-type (WT) and catalytically deficient (dCD) mouse MLL3 and MLL4 proteins (top) and experimental design to explore the changes of histone modifications, gene regulation, and enhancer-promoter (E-P) contacts during neural differentiation in the presence and absence of MLL3/4 catalytic activity (bottom). Tyrosine (Y) residues are mutated to alanine (A) in SET domain to inactivate MLL3/4 in dCD cells. **(B)(C)** Heatmaps showing H3K4me1 (left) and H3K27ac (middle) ChIP-seq and ATAC-seq (right) signals centered at H3K27ac peaks of candidate enhancers identified in WT NPCs. Candidate enhancers were classified based on whether H3K4me1 peak signals were significantly gained only in NPCs (day 5) (*de novo* enhancers, N=3,373) (B) or not (persistent enhancers, N=30,404) (FDR < 0.05, log_2_FC > 0.5) (C), and further classified into enhancers that had significantly lower level of H3K4me1 signals in MLL3/4 dCD NPCs compared with that in WT NPCs (MLL3/4-dependent, N=3,028 and 4,150) and the other enhancers (MLL3/4-independent, N=345 and 26,254). **(D)** The top 5 enriched known TF binding motifs at MLL3/4-dependent (left) and -independent enhancers (right) in the group of *de novo* enhancers. Their enrichment p values and p values of differential motif analysis between MLL3/4-dependent and -independent enhancers are also indicated (see Figure S1I for the same analysis in the group of persistent candidate enhancers). See also Figure S1.

## RESULTS

### MLL3/4 Catalytic Activity Dependent and Independent H3K4me1 at Enhancers during Neural Precursor Cell (NPC) Differentiation

We first analyzed how catalytic inactivation of MLL3/4 methyltransferase altered histone modification genome-wide in MLL3/4 dCD cells during ESC differentiation to NPC. Consistent with previous reports (Dorighi et al., 2017; Hu et al., 2013; Lee et al., 2013; Wang et al., 2016), we observed reduction of H3K4me1 at 19,454 and 25,271 distal elements in ESCs and NPCs, respectively (FDR < 0.05, log_2_ FC > 0.5) (Figures S1A–S1F). H3K27ac signals at the same regions were partially reduced and the degree of changes was positively correlated with the change in H3K4me1 (Figures S1G). Surprisingly, significantly elevated H3K4me1 signals were also observed around a large number of gene promoters (11,001 and 16,175 loci in ESCs and NPCs, respectively), where MLL3/4 occupancy is not detected (Dorighi et al., 2017; Hu et al., 2013; Lee et al., 2013; Wang et al., 2016) (Figures S1B, S1D, and S1F), possibly due to activities of other methyltransferases that bind around promoter regions (Hu et al., 2017; Hu et al., 2013; Hyun et al., 2017). We next focused on the candidate distal enhancers associated with both H3K27ac and H3K4me1 (distance to transcription start site ≥ 10 kb) and measured their dynamic chromatin states by ChIP-seq upon neural precursor differentiation in wild-type (WT) and MLL3/4 dCD cells. In total, 35,744 and 33,777 candidate enhancers were identified in ESCs and NPCs, respectively. During NPC differentiation, 3,373 candidate enhancers gained H3K4me1 signals (FDR < 0.05, log_2_ FC > 0.5) along with increased chromatin accessibility as profiled previously by ATAC-seq (Duren et al., 2017; Xu et al., 2017). Interestingly, 90% (N = 3,028) of them failed to acquire H3K4me1 in MLL3/4 dCD cells, suggesting that MLL3/4 catalytic activity plays a key role in the deposition of H3K4me1 at these *de novo* candidate enhancers during ESC differentiation. Similarly, acquisition of H3K27ac at these distal elements in NPC was also severely impaired in the dCD cells (Figure 1B, Table S2). On the other hand, H3K4me1 level was unaffected at over 26,000 candidate enhancers in MLL3/4 dCD NPCs (Figure 1C). These MLL3/4-independent candidate enhancers were already associated with H3K4me1 in ESC in general and persisted during NPC differentiation. Many of them (14,778 loci) were also annotated as poised enhancers in ESCs and gained H3K27ac signals during NPC differentiation, consistent with a recent report (Dorighi et al., 2017). Interestingly, in the MLL3/4*-*dependent *de novo* candidate enhancers, motifs of GATA family TFs that are known to be important for ESC differentiation and embryonic development were highly enriched (Figure 1D, Figures. S1H and S1I), suggesting a potential role for GATA family TFs in the activation and recruitment of MLL3/4 at these distal enhancers during ESC differentiation (Fujikura et al., 2002; Jozwik et al., 2016; Tremblay et al., 2018; Wamaitha et al., 2015; Yu et al., 2019). These findings suggest that MLL3/4 catalytic activity plays a crucial role in chromatin state of a subset of candidate enhancers, especially those that gained H3K4me1 during NPC differentiation, but is dispensable for the maintenance of H3K4me1 at most candidate enhancers in NPCs. This result highlights both MLL3/4-dependent and -independent mechanisms responsible for H3K4me1 histone modification at enhancers.

### Catalytic Activity of MLL3/4 is Required for Newly Formed E-P Contacts upon NPC Differentiation

We next investigated the changes of chromatin contacts between candidate enhancers and promoters upon loss of MLL3/4 catalytic activity. We performed PLAC-seq (Fang et al., 2016; Mumbach et al., 2016) using antibodies against the promoter histone mark H3K4me3 to map chromatin contacts anchored at active or poised promoters. Previous studies showed that H3K4me3 is not affected by loss of MLL3/4 (Dorighi et al., 2017; Lee et al., 2013). We obtained between 280 and 430 million paired-end reads for each replicate (Table S1). To determine the differential chromatin contacts between compared samples, gene promoters with significantly altered levels of H3K4me3 ChIP-seq signals (DEseq2 (Love et al., 2014), p value < 0.01) were filtered out to remove the antibody bias. In the differential analysis between WT and MLL3/4 dCD cells in ESCs and NPCs, we analyzed about 12,000 gene promoters with similar levels of H3K4me3 ChIP-seq signals using a negative binomial model for each distance-stratified 10-kb interval (Kubo et al., 2021; Li et al., 2014; Su et al., 2019) (Figure S2, Methods). These chromatin contacts analysis showed that the decreases of E-P contacts upon loss of MLL3/4 catalytic activity were predominantly observed in the NPCs (2252 reduced contacts, FDR < 0.05), while much fewer changes were observed in the ESCs (43 reduced contacts, FDR < 0.05) (Figure 2A, Table S3). Meanwhile, the comparison between ESCs and NPCs revealed that the majority of E-P contacts induced during NPC differentiation in WT cells (2545 induced contacts, FDR < 0.05) failed to form in MLL3/4 dCD cells (529 induced contacts, FDR < 0.05) (Figures S3A–S3C). We also identified significant chromatin loops formed between enhancers and promoters using the MAPS algorithm (Juric et al., 2019), and 882 significant E-P contacts (FDR < 0.01) could be detected at the 3,028 MLL3/4-dependent *de novo* enhancers. As expected, they generally displayed increased E-P contacts upon NPC differentiation in WT cells, but such induced E-P contacts were largely diminished in MLL3/4 dCD cells (69 vs 17 of significantly induced contacts, FDR < 0.05), suggesting that the MLL3/4-dependent H3K4me1 is required for the formation of E-P contacts at these putative enhancers during NPC differentiation (Figure 2B). On the other hand, the induction of E-P contacts at the 345 MLL3/4-independent *de novo* enhancers upon NPC differentiation was less affected by the loss of MLL3/4 catalytic activity than at the 3,028 MLL3/4-dependent enhancers (Figures 2C, 2D, and S3D). These results support a general role of MLL3/4-dependent H3K4me1 at enhancers in the establishment of chromatin contacts, as we previously reported (Yan et al., 2018). More importantly, we defined the set of candidate enhancers that are dependent on MLL3/4 catalytic activity for H3K4me1 deposition and formation of long-range E-P contacts during cell differentiation.

**Figure 2.**
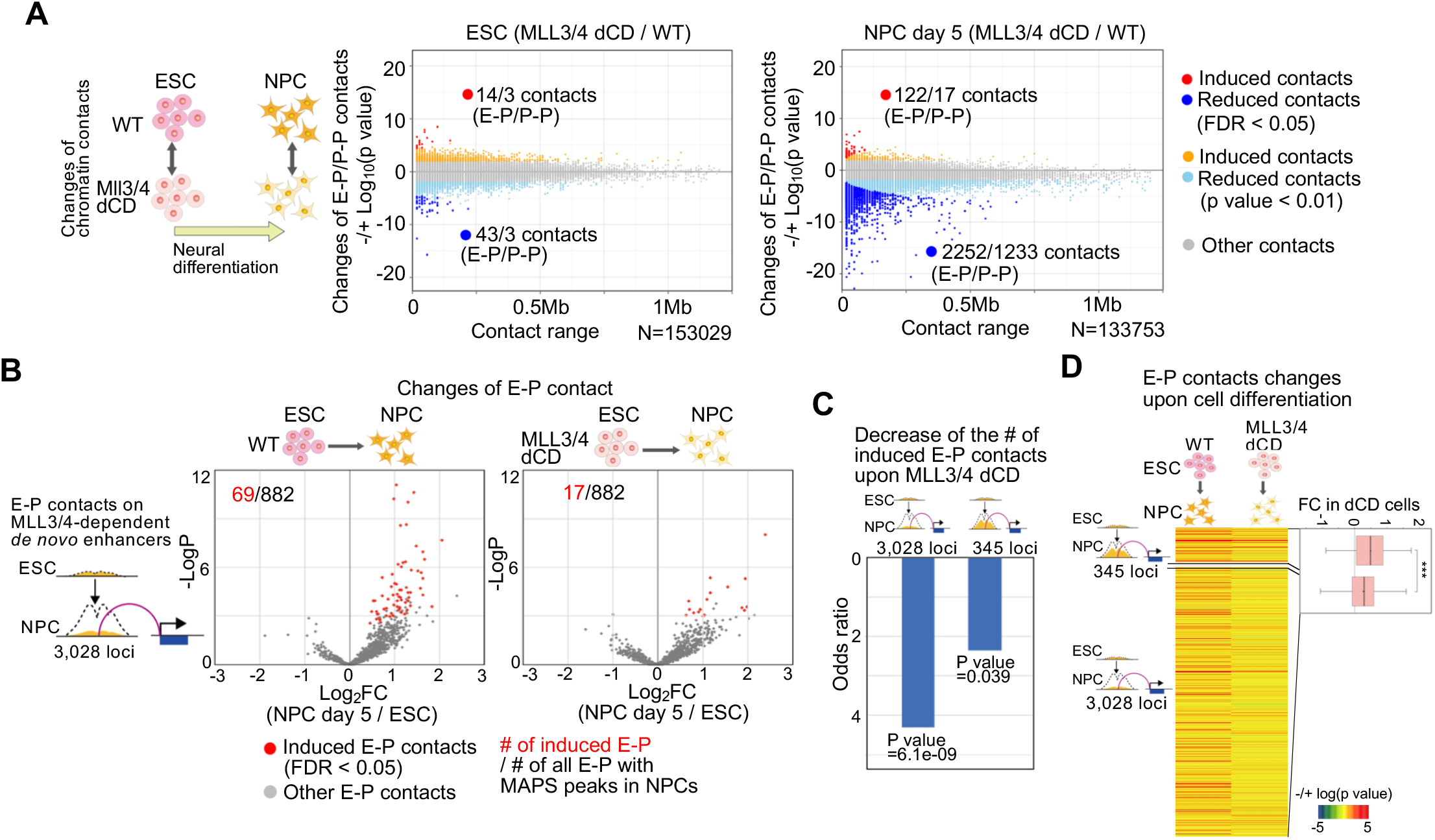
Severe disruption of newly formed E-P contacts during NPC differentiation in MLL3/4 dCD cells. **(A)** Scatter plots showing genome-wide changes of chromatin contacts anchored on promoters and enhancers (y-axis) identified in differential interaction analysis between WT cells and MLL3/4 dCD cells in ESCs (left) and NPCs (right). Genomic distances between their two loop anchor sites are plotted on *x*-axis. The interaction changes are indicated by significance value (−/+log_10_(p-value)). The numbers of significantly changed E-P and promoter-promoter (P-P) contacts are also indicated (FDR < 0.05). Red and orange dots; induced chromatin contacts in MLL3/4 dCD cells (FDR < 0.05 and p value < 0.01, respectively). Blue and light-blue dots; reduced chromatin contacts in MLL3/4 dCD cells (FDR < 0.05 and p value < 0.01, respectively). **(B)** Volcano plots showing changes of E-P contacts centered at the MLL3/4-dependent *de novo* enhancers (3028 loci) upon cell differentiation from ESCs towards NPCs in WT (left) and MLL3/4 dCD cells (right). E-P contacts that were overlapped with significant peaks called by MAPS are plotted (FDR < 0.01). Significantly induced E-P contacts upon cell differentiation (FDR < 0.05) are plotted as red dots and the numbers of them are also indicated. **(C)** Histogram showing the odds ratio of the decrease of the number of significantly induced E-P contacts upon loss of MLL3/4 catalytic activity. E-P contacts centered at the MLL3/4-dependent *de novo* enhancers (3028 loci) and the MLL3/4-independent *de novo* enhancers (345 loci) are shown separately. P values for each odds ratio (Fisher’s exact test) are also indicated. **(D)** Heatmaps showing the changes of E-P contacts centered at the MLL3/4-dependent *de novo* enhancers (3028 loci) and the MLL3/4-independent *de novo* enhancers (345 loci) upon cell differentiation from ESCs towards NPCs in WT (left column) and MLL3/4 dCD cells (right column). The interaction changes are indicated by significance value (−/+log_10_(p-value)). Boxplots also show the changes of E-P contacts by fold change (NPC/WT in MLL3/4 dCD). Central bar, median; lower and upper box limits, 25th and 75th percentiles, respectively; whiskers, minimum and maximum value within the range of (1st quartile-1.5*(3rd quartile-1st quartile)) to (3rd quartile+1.5*(3rd quartile-1st quartile)). *** p value < 0.001, two-tailed t-test. See also Figures S2 and S3.

Meanwhile, we also observed loss of large numbers of promoter-promoter (P-P) contacts in MLL3/4 dCD cells (1,233 contacts in NPCs, FDR < 0.05) (Figure 2A). Interestingly, the changes in P-P contacts upon loss of MLL3/4 catalytic activity did not show any correlation with the changes of H3K4me1 signals, unlike the changes of E-P contacts. (Figure S3E). Instead, the changes of P-P contacts were correlated with the changes of E-P contacts that shared their anchor sites at promoters, suggesting that these P-P contacts might be a consequence of the changes of their nearby E-P contacts (Figures S3F and S3G). Taken together, loss of MLL3/4 catalytic activity caused the failure of H3K4me1 acquisition especially at *de novo* distal enhancers in NPCs, resulting in global disruption of newly formed E-P contacts during neural differentiation.

### Loss of MLL3/4 Catalytic Activity Delays NPC differentiation and Impairs Activation of a Small Number of Genes

We next investigated the impact of the loss of MLL3/4 catalytic activity on gene activation during neural differentiation. The MLL3/4 dCD cells exhibited a delay in formation of neuronal axons and remained in ESC-like round shape colonies after 5 days of the neural induction (Figure 3A). Consistent with this observation, the overall gene expression profiles at each time point also suggested a delay of transcriptional transition in MLL3/4 dCD cells (Figure 3B). While the up-regulation of most NPC marker genes such as *Pax6*, *Sox3*, *Map2, and* the down-regulation of pluripotent markers such as *Pou5f1*, *Sox2* were not interrupted in MLL3/4 dCD cells, induction of other NPC marker genes such as *Tuj1*, *NeuN*, and *Olig2* was significantly delayed (Figure S4A). As a matter of fact, of the 1,303 genes activated during the cell differentiation (FDR < 0.05, FC > 2, RPKM in NPCs > 1), 31.6% or 411 genes (FC > 1.5, FDR < 0.05) were not fully induced in MLL3/4 dCD cells (Figures 3C,S4B–S4D, Table S4). Genes related to organism development and cell differentiation (e.g. Sox11, *Hoxb9*, *Lhx1*) were highly enriched in these genes, suggesting that the loss of MLL3/4 catalytic activity impairs NPC differentiation (Figure 3D).

**Figure 3.**
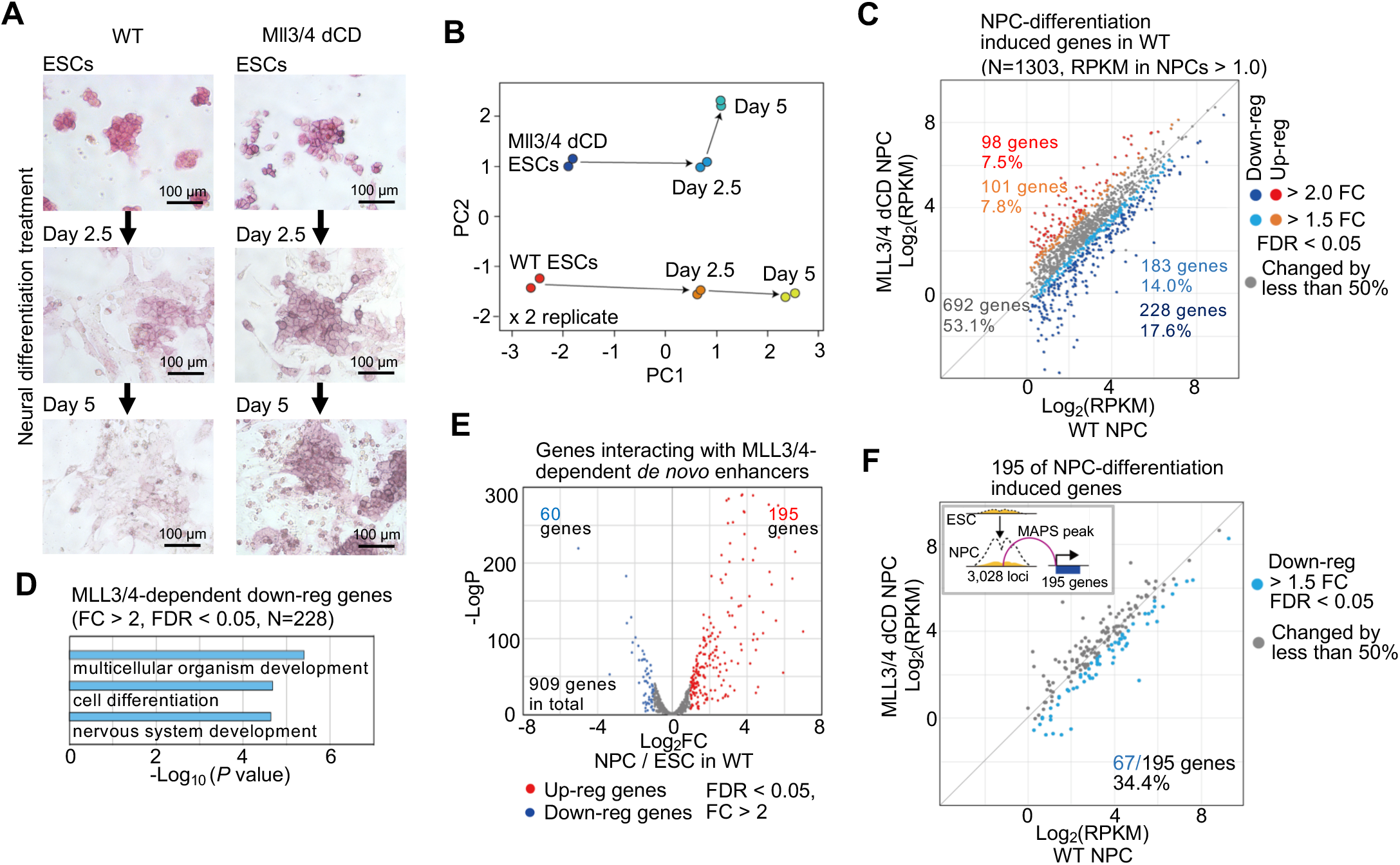
Over 60% of genes interacting with MLL3/4-dependent *de novo* enhancers are still fully induced in MLL3/4 dCD cells during NPC differentiation. **(A)** Microscopic images of cell differentiation from mouse ESCs towards NPCs (day 2.5, 5) in WT and MLL3/4 dCD cells. Alkaline phosphatase staining was performed at each time point. **(B)** Principal component analysis of overall gene expression profiles of WT and Mll3/4 dCD cells at each time point of cell differentiation. Two replicates of each sample are shown. **(C)** Scatter plots showing gene expression levels of NPC-differentiation induced genes (FDR < 0.05, FC > 2, RPKM in NPCs > 1.0) in WT NPCs (x-axis) and MLL3/4 dCD NPCs (day 5) (y-axis). Blue and light-blue dots; down-regulated in MLL3/4 dCD NPCs (FC > 2 and FC > 1.5, respectively, FDR < 0.05). Red and orange dots; up-regulated in MLL3/4 dCD NPCs (FC > 2 and FC > 1.5, respectively, FDR < 0.05). **(D)** Top three enriched GO terms in genes that failed to be up-regulated upon loss of MLL3/4 catalytic activity (228 genes, FC (WT/dCD) > 2, FDR < 0.05 in NPCs). p values (Fisher’s exact test) are also indicated. **(E)** Volcano plots showing gene expression changes between WT ESCs and NPCs in genes that have significant PLAC-seq chromatin contacts (MAPS, FDR < 0.01) with the 3028 of *de novo* MLL3/4-dependent candidate enhancers. Significantly up-regulated and down-regulated genes were plotted as red and blue dots, respectively (FDR < 0.05, FC > 2). **(F)** Scatter plots showing gene expression levels of NPC-differentiation induced genes that had significant interaction with the 3028 of MLL3/4-dependent *de novo* enhancers in WT NPCs (x-axis) and MLL3/4 dCD NPCs (day 5) (y-axis). Down-regulated genes in MLL3/4 dCD NPCs (FC > 1.5, FDR < 0.05) are plotted as light-blue dots. See also Figure S4.

We focused on the induced genes that also displayed chromatin interactions between their promoters and the 3,028 MLL3/4-dependent *de novo* enhancers. In general, genes interacting with these *de novo* candidate enhancers tended to be up-regulated upon cell differentiation in WT cells (Figure 3E). However, over 60% of them continued to be induced in MLL3/4 dCD cells during NPC differentiation despite the reduction of H3K4me1 at distal elements (Figures 3F and S4E). Why do some genes depend on MLL3/4 catalytic activity while others don’t? We hypothesized that genes could be activated by both MLL3/4-dependent and -independent enhancers during cell differentiation (Kubo et al., 2021; Lagha et al., 2012), and the relative fraction of MLL3/4-dependent and -independent enhancers that contact promoters may determine their dependence on the MLL3/4 catalytic activity. To test this hypothesis, we analyzed the chromatin contact counts on two groups of candidate enhancers classified based on the dependence of H3K4me1 on MLL3/4 catalytic activity as defined above (Figures 1B, 1C, and 4A). For each gene, we calculated the ratio of total contact counts on the MLL3/4-independent enhancers to that on all candidate enhancers. Supporting our hypothesis, genes with relatively higher input from MLL3/4-independent enhancers tended to be unaffected by the loss of MLL3/4 catalytic activity than genes making more contacts with MLL3/4-dependent candidate enhancers (Figure 4B). The MLL3/4-independent genes (692 genes, defined in Figure 3C) were more likely to interact with MLL3/4-independent candidate enhancers than the MLL3/4-dependent genes (228 down-regulated and 98 up-regulated genes, defined in Figure 3C) (Figures 4C, S5). These findings suggest that the chromatin contacts with multiple MLL3/4-independent enhancers could sustain the gene activation in the majority of genes during neural differentiation in the absence of MLL3/4 catalytic activity.

**Figure 4.**
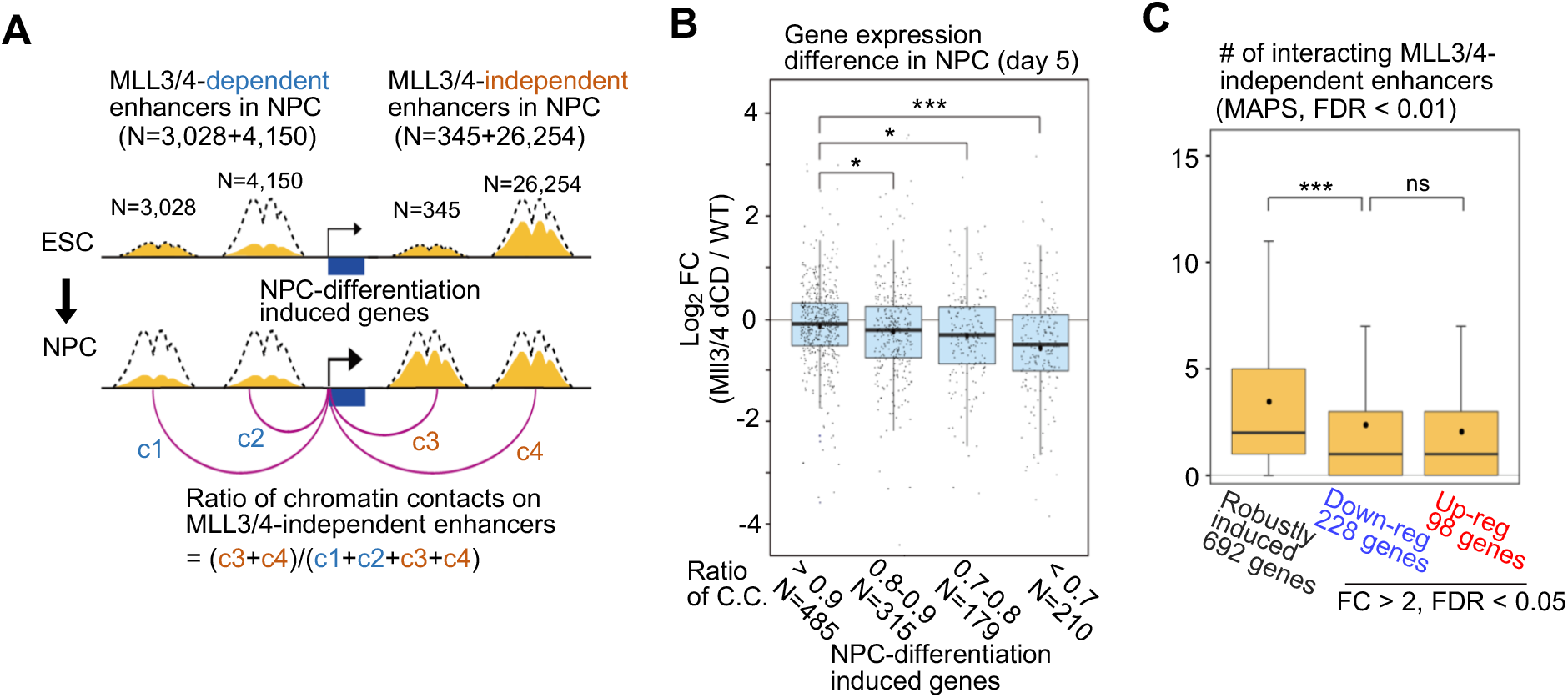
Effects of MLL3/4 catalytic activity loss on enhancer-dependent gene activation during NPC differentiation vary depending on the relative promoter contacts from MLL3/4- dependent candidate enhancers. **(A)** Schematic representation of the method of calculating the ratio of chromatin contacts between promoters and MLL3/4-dependent or independent candidate enhancers. Total contact counts on a gene promoter and multiple MLL3/4-independent enhancers are divided by total contact counts on a gene promoter and all candidate enhancers in WT NPCs (contact range ≥ 10 kb). See Methods for details of the calculation. **(B)** Boxplots showing changes in gene expression between WT and MLL3/4 dCD cells during NPC differentiation. NPC-differentiation induced genes were classified into four groups based on the ratios of chromatin contact between promoters and MLL3/4-independent enhancers versus all chromatin contacts anchored at the same promoters. The ratios and the numbers of genes are indicated on the bottom. Central bar, median; lower and upper box limits, 25th and 75th percentiles, respectively; whiskers, minimum and maximum value within the range of (1st quartile-1.5*(3rd quartile- 1st quartile)) to (3rd quartile+1.5*(3rd quartile- 1st quartile)). * p value < 0.05, *** p value < 0.001, one-tailed t-test. **(C)** Boxplots showing the number of MLL3/4-independent candidate enhancers with significant chromatin contacts as determined by PLAC-seq assays (MAPS, FDR < 0.01) with each group of genes. These NPC-differentiation induced genes were classified based on the differential gene expression analysis in Figure 3C. Central bar, median; lower and upper box limits, 25th and 75th percentiles, respectively; whiskers, minimum and maximum value within the range of (1st quartile- 1.5*(3rd quartile- 1st quartile)) to (3rd quartile+1.5*(3rd quartile- 1st quartile)). ns p value > 0.05, *** p value < 0.001, one-tailed t-test. See also Figure S5.

## DISCUSSION

In summary, our study found that accumulation of H3K4me1 and H3K27ac at new candidate enhancers established during NPC differentiation were generally dependent on catalytic activities of MLL3 and MLL4. By contrast, H3K4me1 at a much larger number of candidate enhancers is independent of Mll3/4 catalytic activity and could be catalyzed by other histone methyltransferases. Furthermore, consistent with our previous reports (Dixon et al., 2015; Yan et al., 2018), we observed a severe disruption of newly formed E-P contacts at these new candidate enhancers in MLL3/4 dCD NPCs. Lastly, our study demonstrated that loss of MLL3/4 catalytic activity delays NPC differentiation and activation of nearly 1/3 of lineage specific genes, and our results support that MLL3/4-dependent H3K4me1 plays a significant role at E-P contacts at candidate enhancers which in turn contributes to gene activation at a subset of genes that make more contacts with them. Notably, although MLL3/4 are believed to be the major regulators of H3K4me1 in mammalian cells, H3K4me1 at over 75% of candidate distal enhancers in NPCs was independent of MLL3/4 catalytic activity. Future studies are required to assess whether this finding can be generalized to other cell lineages, and to determine the histone methyltransferase that mediate the H3K4me1 at these elements (Crump and Milne, 2019; Hyun et al., 2017). Additionally, further exploration of TFs such as GATA family members as shown in this study would help to unveil the cell type specific role of MLL3/4 (Jozwik et al., 2016). A recent preprint study also focused on the role of MLL3/4 catalytic activity in mouse embryonic development and ESC differentiation and reported their altered transcription in subsets of genes (Xie et al., 2020). Although the impact of MLL3/4 catalytic activity loss on those gene regulation programs apparently differs depending on tissue types and differentiation conditions, our study provides an explanation of the observed MLL3/4-dependent and -independent gene activations by focusing on the E-P contact dynamics. It should be noted that our analysis of E-P contacts does not include the information of chromatin contacts within 10-kb genomic distance due to a limitation of resolution in the current approach. Our analysis is also not based on acute depletion of Mll3/4 catalytic activity, and some genes might be affected by secondary effects of the long-term culturing. Nevertheless, our findings clarify the functional role of MLL3/4 catalytic activity in gene regulation programs and provide new insight into the role of MLL3/4-mediated H3K4me1 in cell differentiation.

## METHODS

### Contact for Reagent and Resource Sharing

Further information and requests for reagents may be directed to and will be fulfilled by the corresponding author Bing Ren and the first author Naoki Kubo.

### EXPERIMENTAL MODEL AND SUBJECT DETAILS

#### Cell lines

Mouse R1 ES cell line was used for MLL3/4 catalytically deficient ES cell line, which was reported in the previous study (Dorighi et al., 2017). ESCs were cultured in KnockOut Serum Replacement containing mouse ES cell media: DMEM 85%, 15% KnockOut Serum Replacement (Gibco), penicillin/streptomycin (Gibco), 1× non-essential amino acids (Gibco), 1× GlutaMax (Gibco), 1000 U/ml LIF (Millipore), 0.4 mM β-mercaptoethanol. The cells were grown on 0.2% gelatin-coated plates with irradiated mouse embryonic fibroblasts (MEFs) (GlobalStem). Cells were maintained by passaging using Accutase (Innovative Cell Technologies) at 37°C and 5% CO2. Medium was changed daily when cells were not passaged. Cells were checked for mycoplasma infection and tested negative.

### METHOD DETAILS

#### Neural progenitor cell differentiation

NPC differentiation was conducted utilizing retinoic acid (RA) induction (Bain et al., 1995; Strubing et al., 1995). ESCs were grown on MEFs and passaged on 0.2% gelatin-coated plates without MEFs one day before starting differentiation treatment. On day 0, LIF was deprived from the culture medium. From day 1, 5 uM RA (Sigma, R2625) was added with the LIF-deprived medium. Cells were harvested on day 2.5 and day 5. Alkaline phosphatase staining was performed at each time point using the AP Staining kit II (Stemgent, 00-0055).

#### ChIP-seq library preparation

ChIP-seq experiments for each histone mark were performed as described in ENCODE experiment protocols (“Ren Lab ENCODE Chromatin Immunoprecipitation Protocol” in https://www.encodeproject.org/documents/) with minor modifications. Cells were crosslinked with 1% formaldehyde for 10 minutes. We used 1.0 million cells for each ChIP sample. Shearing of chromatin was performed using truChIP Chromatin Shearing Reagent Kit (Covaris) according to the manufacturer’s instructions. Covaris M220 was used for sonication with following parameters: 10 minutes duration at 10.0% duty factor, 75.0 peak power, 200 cycles per burst at 5-9°C temperature range. For immunoprecipitation, we used 11 μL anti-rabbit or anti-mouse IgG Dynabeads (Life Technologies) and wash them with cold BSA/PBS (0.5 mg / mL bovine serum albumin in 1x phosphate buffered saline) for 3 times. After washing, 3 μg antibody with 147 μL cold BSA/PBS were added to the beads and incubated over 2 hours at 4°C. After incubation, beads were washed with150 μL cold BSA/PBS for 3 times and mixed with 100 μL Binding Buffer (1% Triton X-100, 0.1% Sodium Deoxycholate, 1x complete protease inhibitor (Roche)) plus 100 μL 0.2 μg/μl chromatin followed by overnight incubation on a rotating platform at 4°C. Beads were washed 5 times with 50 mM Hepes pH 8.0, 1% NP-40, 1 mM EDTA, 0.70% Sodium Deoxycholate, 0.5 M LiCl, 1x complete protease inhibitor (Roche) and washed once with 150 μL cold 1x TE followed by incubation at 65°C for 20 minutes in 150 μL ChIP elution buffer (10 mM Tris-HCl pH 8.0, 1 mM EDTA, 1% SDS). The beads were removed and the samples were incubated at 65°C overnight to reverse crosslinks. The input samples were also processed in parallel with the ChIP samples. Samples were incubated with RNase A (final conc. = 0.2 mg/mL) at 37°C for 1 hour, and Proteinase K (final conc. = 0.4 mg/mL) at 55°C for 1 hour. The samples were extracted with phenol: chloroform: isoamyl alcohol (25:24:1) and precipitated with ethanol. We used 3-5 ng of starting IP materials for preparing Illumina sequencing libraries. The End-It DNA End-Repair Kit (Epicentre) was used to repair DNA fragments to blunt ends. A-tailing 3’ end was performed using Klenow Fragment (3’→5’ exo-) (New England Biolabs), and then TruSeq Adapters were ligated by Quick T4 DNA Ligase (New England Biolabs). Size selection using AMpure Beads (Beckman Coulter) was performed to get 300-500bp DNA and PCR amplification (8-10 cycles) was performed. Libraries were sequenced on HiSeq4000 single end for 50 bp. Two biological replicates were prepared for each sample.

#### RNA-seq library preparation

Total RNA was extracted using the AllPrep Mini kit (QIAGEN) according to the manufacturer’s instructions and 1 μg of total RNA was used to prepare each RNA-seq library. The libraries were prepared using TruSeq Stranded mRNA Library Prep Kit (Illumina). Libraries were sequenced on HiSeq4000 using 50 bp paired-end. Two biological replicates were prepared for each sample.

#### PLAC-seq library preparation

PLAC-seq experiments were performed as previously described (Fang et al., 2016). Cells were crosslinked with 1% formaldehyde (w/v, methanol-free, ThermoFisher) for 15 minutes. The crosslinked pellets (2.5–3 million cells per sample) were incubated with 300ul of lysis buffer (10 mM Tris-HCl pH 8.0, 10 mM NaCl, 0.2% Igepal CA630, 33 μL, 1x complete protease inhibitor (Roche)) on ice for 15 min, washed with 500 μL cold lysis buffer, and then incubated in 50uL of 0.5% SDS for 10 min at 62°C. After heating, 160 μL of 1.56% Triton X-100 was added and incubated for 15min at 37°C. To digest chromatin 100U MboI and 25uL of 10X NEBuffer2 were added followed by 2 hours incubation at 37°C with agitation at 900rpm. MboI was inactivated by heating at 62°C. Digestion efficiency was confirmed by performing agarose gel electrophoresis of the samples. The digested ends were labeled with biotin by adding 37.5uL of 0.4mM biotin-14-dATP (Life Tech), 1.5 μL of 10mM dCTP, 10mM dTTP, 10mM dGTP, and 8uL of 5U/ul Klenow (New England Biolabs) and incubating at 37°C for 1 hour with shaking at 900 rpm. Then the samples were mixed with 1x T4 DNA ligase buffer (New England Biolabs), 0.83% Trition X-100, 0.1 mg/mL BSA, 2000U T4 DNA Ligase (New England Biolabs, M0202), and incubated at room temperature for 2 hours with shaking with slow rotation. The ligated cell pellets were resuspended in 125 ul of RIPA buffer with protease inhibitor and incubated on ice for 10 minutes. The cell lysates were sonicated using Covaris M220. After spinning, we saved 20 ul supernatant as input, and for the rest part, 100 ul of antibody-coupled beads were added to the supernatant sample, and then rotated in cold room at least 12 hours. For immunoprecipitation, 300 ul of M280 sheep anti-rabbit IgG beads (ThermoFisher) was washed with cold BSA/PBS (0.5 mg / mL bovine serum albumin in 1x phosphate buffered saline) for 4 times. After washing, 30 ug anti-H3K4me3 (Millipore, 04-745) with 1 mL cold BSA/PBS were added to the beads and incubated on a rotating platform at 4°C for over 3 hours. After incubation, beads were washed with cold BSA/PBS and resuspended in 600 ul RIPA buffer. The beads were washed with RIPA buffer (3 times), RIPA buffer + 0.16M NaCl (2 times), LiCl buffer (1 time), and TE buffer (2 times) at 4°C for 3 minutes at 1000 rpm. For reverse crosslinking, 163 ul extraction buffer (135 ul 1xTE, 15 ul 10% SDS, 12 ul 5M NaCl, 1 ul RnaseA (10mg/ml)) was added and incubated at 37°C for 1 hour at 1000 rpm, and 20 ug of proteinase K was added and incubated at 65°C for 2 hours at 1000rpm. After crosslinking, DNA was purified using Zymo DNA clean & concentrator and eluted with 50 ul of 10mM Tris (pH 8.0). For biotin enrichment, 25 ul of T1 Streptavidin Beads (Invitrogen) per sample were washed with 400 ul Tween wash buffer (5 mM Tris-HCl pH 8.0, 0.5 mM EDTA, 1 M NaCl, 0.05% Tween-20), and resuspended in 50 ul of 2x Binding buffer (10 mM Tris-HCl pH 7.5, 1 mM EDTA, 2 M NaCl). The purified 50 ul DNA sample was added to the 50 ul resuspended beads and incubated at room temperature for 15 minutes with rotation. The beads were washed with 500 ul of Tween wash buffer twice and washed with 100 ul Low EDTA TE (supplied by Swift Biosciences kit). Then beads were resuspended in 40 ul Low EDTA TE. Next, we used Swift Biosciences kit (Cat. No. 21024) for library construction with modified protocol. The Repair I Reaction Mix was added to 40 ul sample beads and incubated at 37°C for 10 minutes at 800 rpm. The beads were washed with 500 ul Tween wash buffer twice and washed with 100 ul Low EDTA TE once. The Repair II Reaction Mix was added to the beads followed by incubation at 20°C for 20 minutes at 800 rpm. The beads were washed in the same way with 500 ul Tween wash buffer and 100 ul Low EDTA TE. Then, 25 ul of the Ligation I Reaction Mix and Reagent Y2 was added to the beads followed by incubation at 25°C for 15 minutes at 800 rpm. The beads were washed with Tween wash buffer and Low EDTA TE. Then 50 ul of the Ligation II Reaction Mix was added to the beads followed by incubation at 40°C for 10 minutes at 800 rpm. The beads were washed and resuspended in 21 ul 10mM Tris-HCl (pH 8.0). The amplification and purification were performed according to the Swift library kit protocols. Libraries were sequenced on Illumina HiSeq 4000. Two biological replicates were prepared for each sample.

### QUANTIFICATION AND STATISTICAL ANALYSIS

#### ChIP-seq data analysis

Each fastq file was mapped to mouse genome (mm10) with BWA(Li and Durbin, 2009) -aln with “-q 5 -l 32 -k 2” options. PCR duplicates were removed using Picard MarkDuplicates (https://github.com/broadinstitute/picard) and the bigWig files were created using deepTools (Ramirez et al., 2016) with following parameters: bamCompare --binSize 10 --normalizeUsing RPKM --operation subtract (or log2). The deepTools was also used for generating heatmaps. Peaks were called with input control using MACS2 (Zhang et al., 2008) with broad peak calling. The candidate active enhancer regions were characterized by the presence of both H3K4me1 and H3K27ac peaks, but not H3K4me3 peaks. DEseq2 (Love et al., 2014) was used for differential peak analysis. We defined “*De novo* enhancers in NPC” as the enhancer regions whose H3K4me1 signal was significantly increased from ESCs to NPCs (FDR < 0.05, log_2_ FC > 0.5). The same differential peak analysis was performed between WT and MLL3/4 dCD cells to determine the MLL3/4-dependent and -independent enhancers.

#### Motif analysis

Enrichment analysis of known DNA binding motifs was performed using HOMER tool (Heinz et al., 2010). Default parameters with a fragment size of 1000 bp and “-mask” parameter were used. In the differential motif analysis, regions from the compared MLL3/4-dependent or - independent enhancers were used as background by adding “-bg” parameter.

#### RNA-seq data analysis

RNA-seq reads (paired-end) were mapped to the mm10 genome using STAR (Dobin et al., 2013). The mapped reads were counted using HTSeq (Anders et al., 2015) and the output files from two replicates were subsequently analyzed by edgeR (Robinson et al., 2010) to detect the differentially expressed genes (FDR < 0.05, FC > 2 or FC > 1.5). RPKM was calculated using an in-house pipeline.

#### ATAC-seq data analysis

ATAC-seq reads (paired-end) were mapped to the mm10 genome and processed using ENCODE ATAC-seq pipeline (https://github.com/ENCODE-DCC/atac-seq-pipeline). The deepTools (Ramirez et al., 2016) was used to generate bigwig files and heatmaps as described above.

#### PLAC-seq data analysis

PLAC-seq reads (paired-end) were aligned against the mm10 genome using BWA -mem (Li and Durbin, 2009). PCR duplicate reads were removed using Picard MarkDuplicates. Filtered reads were binned at 10 kb size to generate the contact matrix. Individual bins that were overlapped with H3K4me3 peaks on transcription start sites (TSSs) were used for downstream differential contact analysis. MAPS (Juric et al., 2019) was used for peak calling with default settings in 10 kb resolution. For the differential contact analysis (Kubo et al., 2021), the raw contact counts in 10 kb resolution bins that have the same genomic distance were used as inputs. To minimize the bias from genomic distance, we stratified the inputs into every 10-kb genomic distance from 10 kb to 150 kb, and the rest of the input bins with longer distances were stratified to have a uniform size of input bins that were equal to that of 140–150 kb distance bins. Since these inputs showed negative binomial distribution, edgeR (Robinson et al., 2010) was used to get the initial set of differential interactions. Only bins that had more than 20 contact counts in each sample of two replicates were used for the downstream analysis. The significances of these differential interactions are either due to the difference in their H3K4me3 ChIP coverage or 3D contacts coverage. Therefore, the chromatin contacts from promoter regions with differential ChIP-seq peaks between the samples (p value < 0.01) were removed and only the chromatin contacts with the same level of H3K4me3 ChIP-seq peaks were processed. We used all bins for inputs that included non-significant interactions that were not identified by MAPS peak caller because the majority of short-range interactions were not identified as significant peaks due to their high background and the changes in these short-range interactions might be also important for gene regulation. We identified a large number of differentially changed short-range interactions even though many of them were not identified by peak caller, and we observed that such differentially changed interactions were positively correlated with the changes of H3K4me1/H3K27ac levels on their anchor sites during neural differentiation (Figure S3C), suggesting these interaction changes might reflect the biological changes. We used significance level with change direction (−/+ log_10_(p-value)) instead of fold change to show the changes of chromatin contacts because fold change tends to be marginal when it is short-range interaction although their changes are actually significant and biologically meaningful. To visualize the chromatin contacts, we used WashU Epigenome Browser (Zhou et al., 2013).

In Figure 4, the ratio of chromatin contacts on MLL3/4-independent enhancers was calculated by simply summing raw contact counts between promoters and all MLL3/4-independent enhancers and dividing by total contact counts between promoters and all candidate enhancers in each gene in WT NPCs. The contact counts within 10-kb distance (1 bin) was not added. Genes that had less than 50 of the total contacts counts were removed for downstream analysis.

#### Odds ratio calculation for CTCF-dependent E-P contacts enrichment

For Figure S5B, all genes were classified based on the distance to the nearest interacting enhancer and the number of enhancers around TSS (< 200 kb) (categorized into 3×3 bins). The distance to the nearest interacting enhancer is represented by the shortest genomic distance of significant PLAC-seq peaks on enhancers and promoters (p-value < 0.01). Then, we generated 2×2 tables based on whether they are stably-regulated genes or not (FDR < 0.05) and whether they were categorized into the bin or not. Odds ratios and p-values on each 2×2 tables were calculated. In Figure S5C and S5D, the same analysis as panel (B) was performed in differentially down-regulated genes and differentially up-regulated genes.

### DATA AND SOFTWARE AVAILABILITY

All datasets generated in this study have been deposited to Gene Expression Omnibus (GEO), with accession number GSE160892. ATAC-seq datasets were downloaded from GSE84646 and GSE98479 (Duren et al., 2017; Xu et al., 2017).

### KEY RESOURCES TABLE

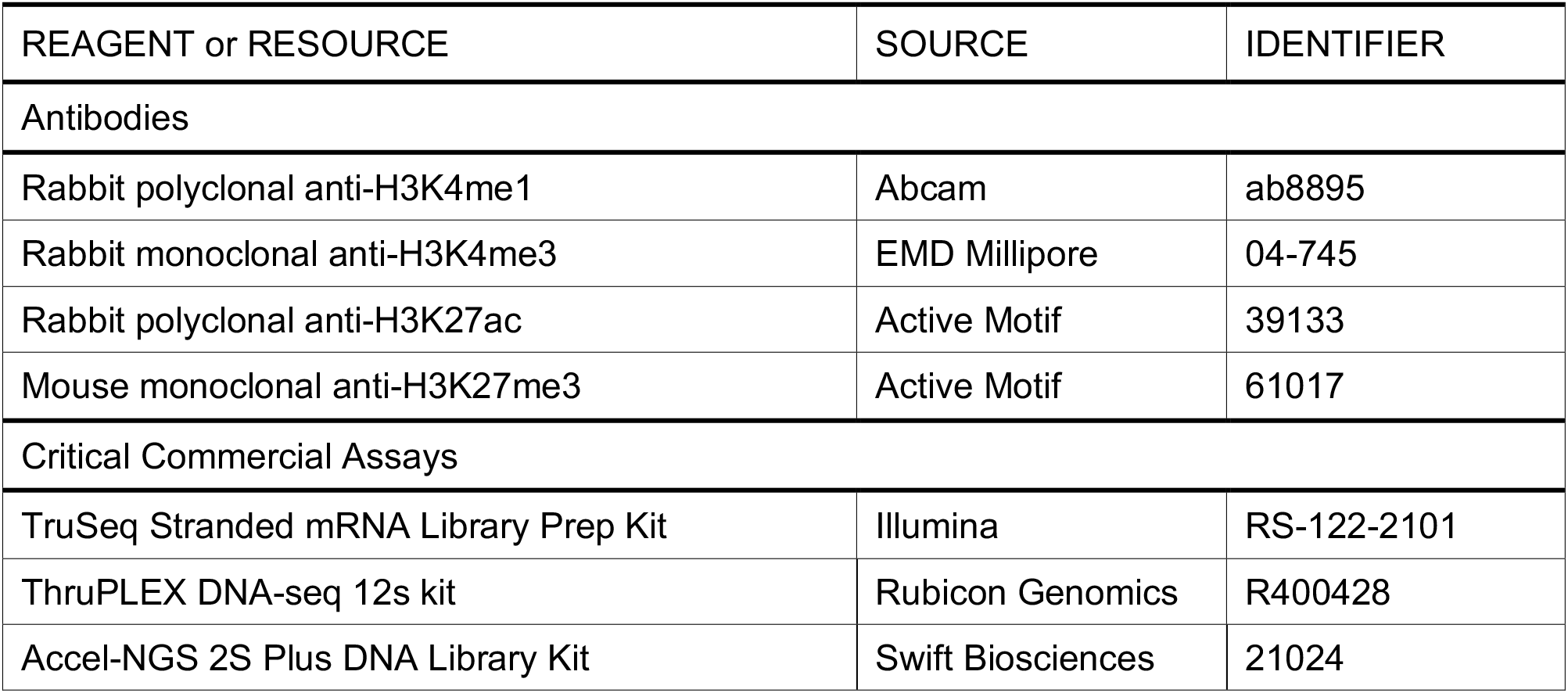
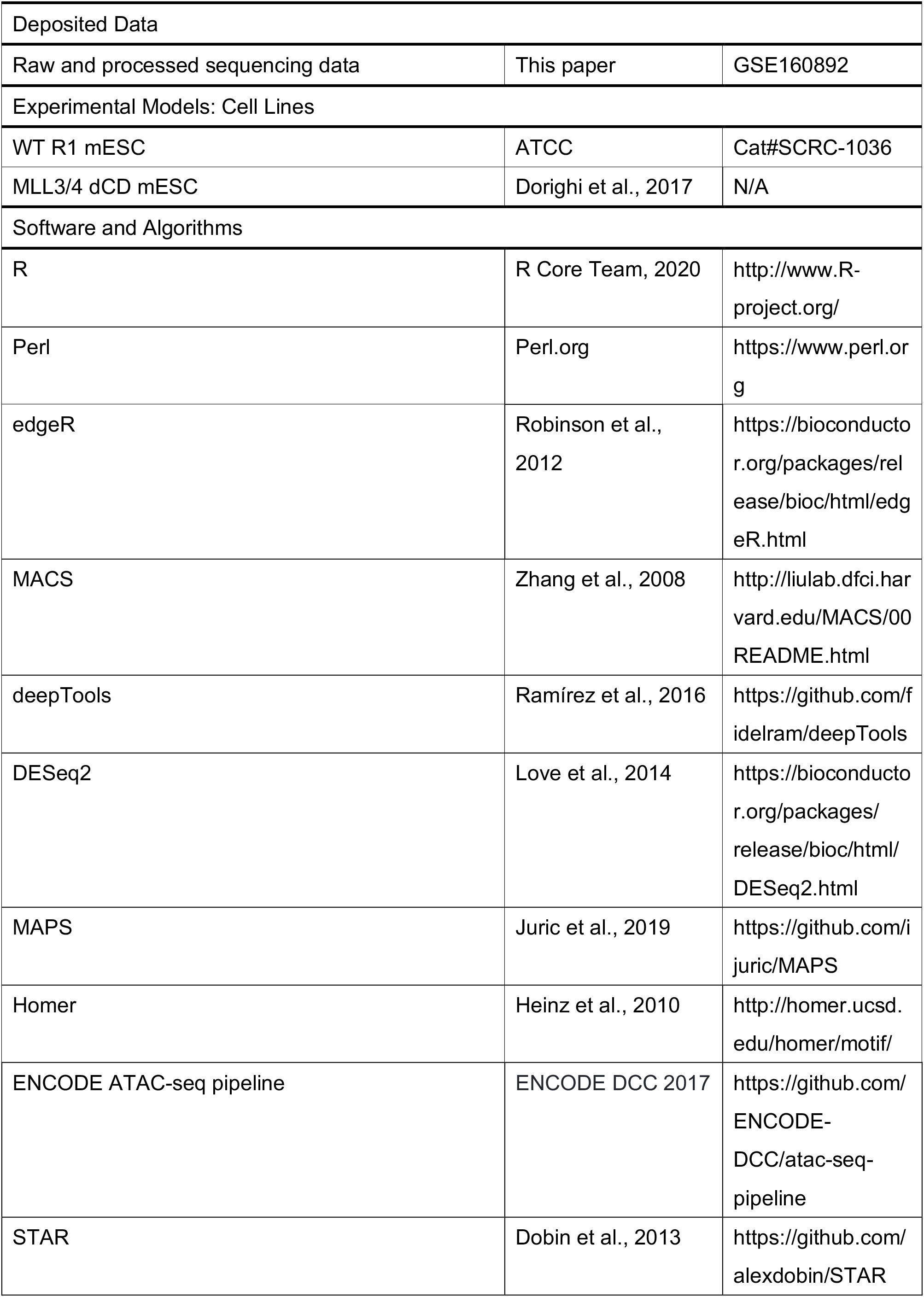
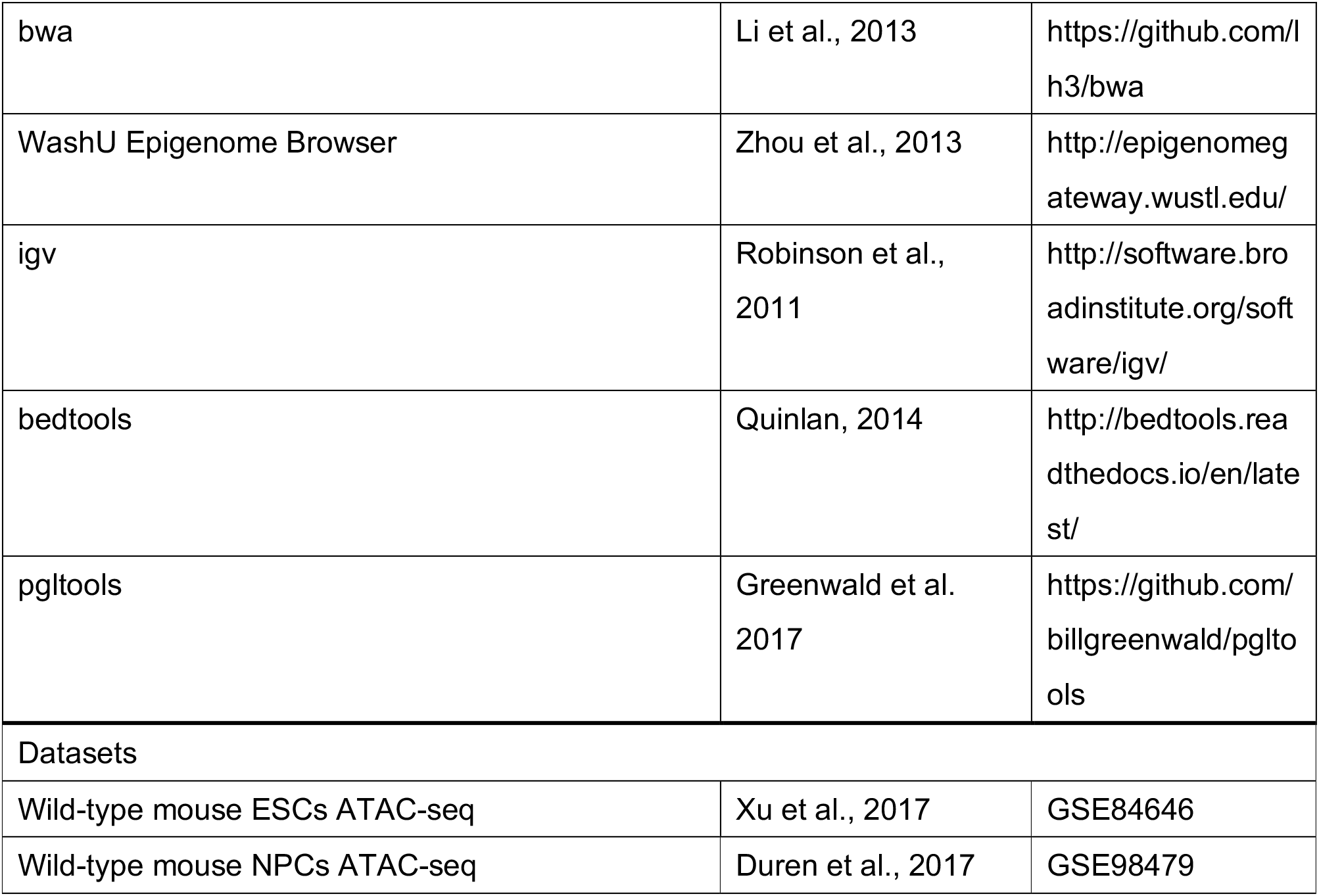

| REAGENT or RESOURCE | SOURCE | IDENTIFIER |
| --- | --- | --- |
| Antibodies |  |  |
| Rabbit polyclonal anti-H3K4me1 | Abcam | ab8895 |
| Rabbit monoclonal anti-H3K4me3 | EMD Millipore | 04-745 |
| Rabbit polyclonal anti-H3K27ac | Active Motif | 39133 |
| Mouse monoclonal anti-H3K27me3 | Active Motif | 61017 |
| Critical Commercial Assays |  |  |
| TruSeq Stranded mRNA Library Prep Kit | Illumina | RS-122-2101 |
| ThruPLEX DNA-seq 12s kit | Rubicon Genomics | R400428 |
| Accel-NGS 2S Plus DNA Library Kit | Swift Biosciences | 21024 |
| Deposited Data |  |  |
| Raw and processed sequencing data | This paper | GSE160892 |
| Experimental Models: Cell Lines |  |  |
| WT R1 mESC | ATCC | Cat#SCRC-1036 |
| MLL3/4 dCD mESC | Dorigi et al., 2017 | N/A |
| Software and Algorithms |  |  |
| R | R Core Team, 2020 | http://www.R-project.org/ |
| Perl | Perl.org | https://www.perl.org |
| edgeR | Robinson et al., 2012 | https://bioconductor.org/packages/release/bioc/html/edgeR.html |
| MACS | Zhang et al., 2008 | http://liulab.dfci.harvard.edu/MACS/00README.html |
| deepTools | Ramírez et al., 2016 | https://github.com/fidelram/deepTools |
| DESeq2 | Love et al., 2014 | https://bioconductor.org/packages/release/bioc/html/DESeq2.html |
| MAPS | Juric et al., 2019 | https://github.com/ijuric/MAPS |
| Homer | Heinz et al., 2010 | http://homer.ucsd.edu/homer/motif/ |
| ENCODE ATAC-seq pipeline | ENCODE DCC 2017 | https://github.com/ENCODE-DCC/atac-seq-pipeline |
| STAR | Dobin et al., 2013 | https://github.com/alexdobin/STAR |
| bwa | Li et al., 2013 | https://github.com/lh3/bwa |
| WashU Epigenome Browser | Zhou et al., 2013 | http://epigenomegateway.wustl.edu/ |
| igv | Robinson et al., 2011 | http://software.broadinstitute.org/software/igv/ |
| bedtools | Quinlan, 2014 | http://bedtools.readthedocs.io/en/latest/ |
| pgltools | Greenwald et al. 2017 | https://github.com/billgreenwald/pgltools |
| Datasets |  |  |
| Wild-type mouse ESCs ATAC-seq | Xu et al., 2017 | GSE84646 |
| Wild-type mouse NPCs ATAC-seq | Duren et al., 2017 | GSE98479 |

## ACKNOWLEDGMENTS

We would like to thank Drs. Kristel M Dorighi and Joanna Wysocka (Stanford School of Medicine) for sharing the MLL3/4 dCD mouse ES cell line. We would like to give special thanks to Samantha Kuan for operating the sequencing instruments and Robert Morey for helping with experiments. We would like to acknowledge the help of Drs. Ivan Juric, Armen Abnousi, and Ming Hu (Lerner Research Institute, Cleveland Clinic Foundation) for sharing computational pipelines. We would also like to give special thanks to Drs. Bin Li, Miao Yu, Ramya Raviram, Yanxiao Zhang, and Yang Li for sharing helpful computational pipelines and protocols, as well as all the other members of the Ren laboratory. This work was supported by the Ludwig Institute for Cancer Research (B.R.), NIH (1U54DK107977-01) (B.R.), and a Postdoc fellowship from the TOYOBO Biotechnology Foundation (N.K.).

## AUTHOR CONTRIBUTIONS

N.K. and B.R. conceived the project. N.K., R.H., and Z.Y. carried out library preparation. N.K. performed data analysis. N.K. and B.R. wrote the manuscript. All authors edited the manuscript.

## Supplementary figure legends

**Figure S1.**
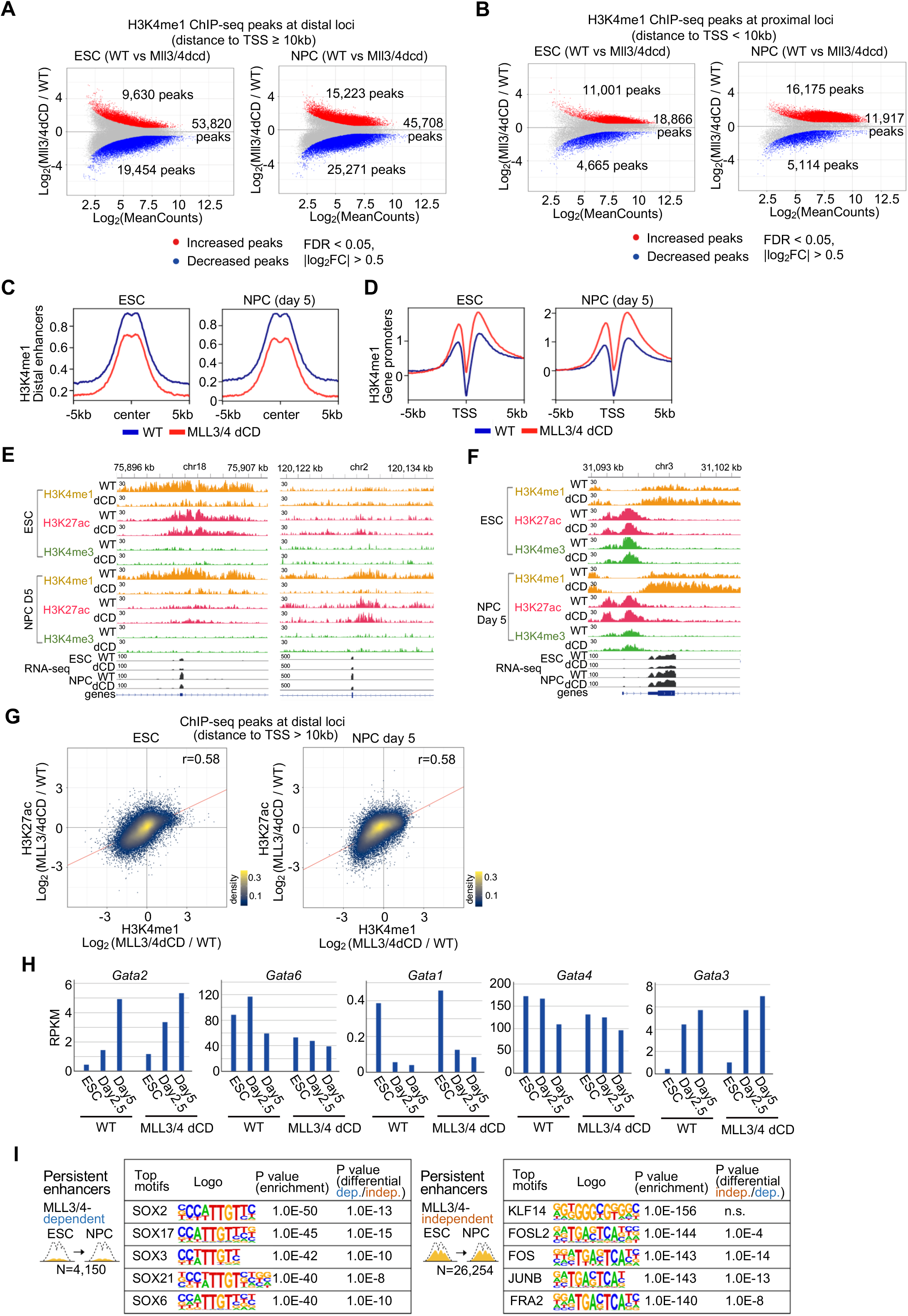
Changes of histone ChIP-seq peak levels between WT and MLL3/4 dCD cells. Related to Figure 1. **(A)(B)** Scatter plots showing changes of H3K4me1 ChIP-seq peak signals between WT and Mll3/4 dCD cells in ESCs (left) and NPCs (day 5) (right). H3K4me1 ChIP-seq peaks at distal loci (distance to transcription start site (TSS) ≥ 10 kb) (A) and peaks at proximal loci (distance to TSS < 10 kb) (B) are plotted separately. Significantly increased and decreased peaks are plotted as red and blue dots, respectively (FDR < 0.05, log_2_FC > 0.5). **(C)(D)** Average enrichments of H3K4me1 ChIP-seq signals around distal enhancers (distance to TSS ≥ 10 kb) (C) and gene promoters (TSSs) (D) in ESCs and NPCs. Blue lines: WT. Red lines: MLL3/4 dCD. **(E)** Genome browser snapshots of regions around a candidate enhancer whose H3K4me1 signal was significantly decreased upon loss of MLL3/4 catalytic activity in ESCs and NPCs (left) and an enhancer whose H3K4me1 was significantly decreased only in NPCs (right). H3K4me1, H3K27ac, and H3K4me3 ChIP-seq and RNA-seq are shown. **(F)** Genome browser snapshots of a region around gene promoter whose H3K4me1 signal was significantly increased upon loss of MLL3/4 catalytic activity. **(G)** Scatter plots showing changes of H3K4me1 peak signals (x-axis) and changes of H3K27ac peak signals (y-axis) upon loss of MLL3/4 catalytic activity in ESCs (left) and NPCs (right). Peaks at distal loci (distance to TSS ≥ 10 kb) are shown. Pearson correlation coefficients and linear trendlines (red line) are also indicated. **(H)** Gene expression profiles of *Gata* family members. RPKM (reads per kilobase of transcript, per million mapped reads) values of RNA-seq (average of two replicates) at each time point in WT and MLL3/4 dCD cells are shown. **(I)** The top 5 enriched known TF binding motifs at MLL3/4-dependent (left) and -independent enhancers (right) in the group of persistent candidate enhancers. Their enrichment p values and p values of differential motif analysis between MLL3/4-dependent and -independent enhancers are also indicated (see Figure 1D for the same analysis in the group of *de novo* enhancers in NPCs).

**Figure S2.**
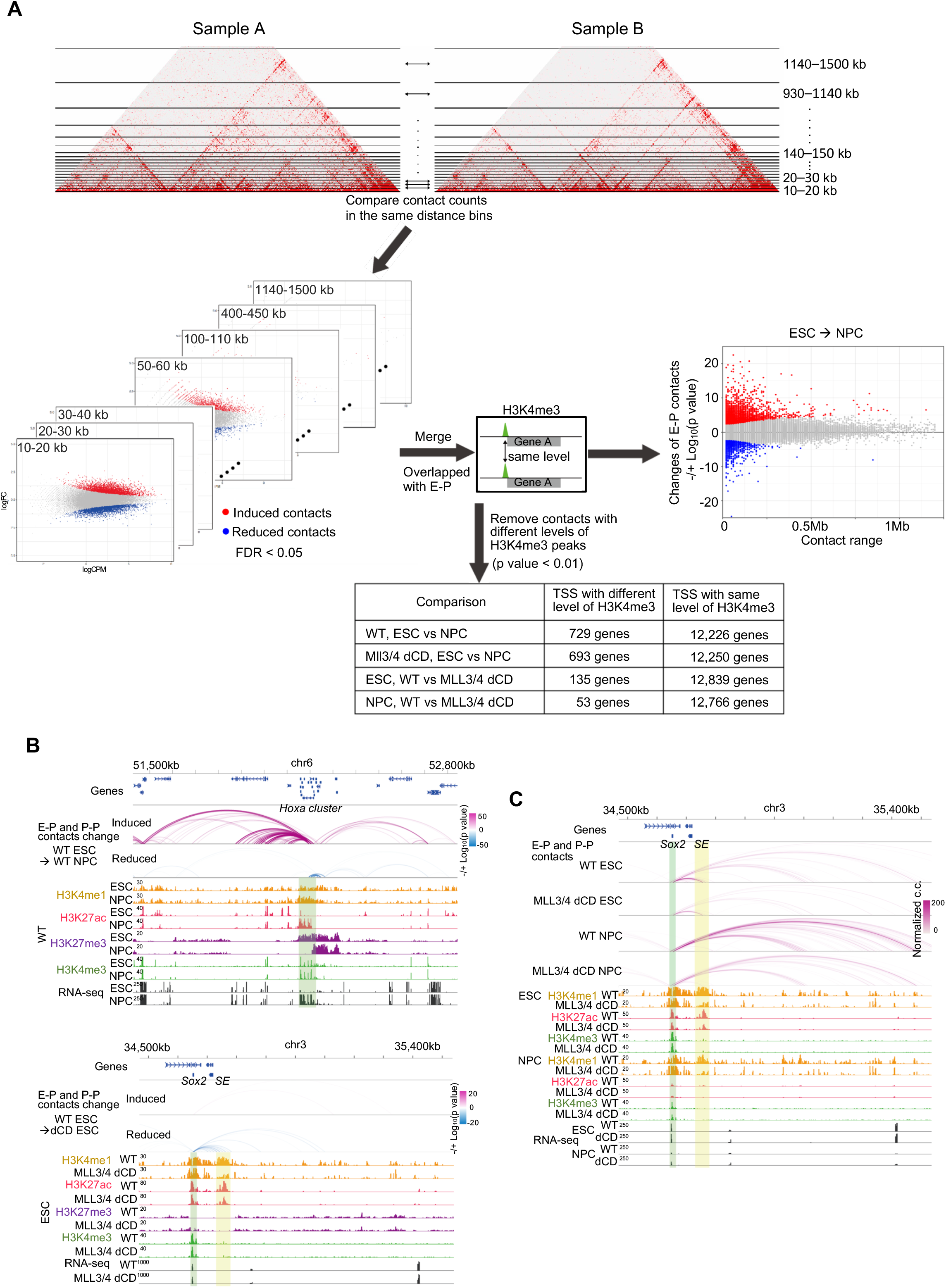
Differential chromatin contact analysis. Related to Figure 2. **(A)** Schematic representation of the method for the differential analysis of H3K4me3 PLAC-seq datasets. The input contact matrix bins were stratified into every 10-kb genomic distance from 10-kb to 150-kb and the rest of the bins of longer distances were stratified to have a uniform number of input bins that were equal to that of 140–150 kb distance bins, and the contact counts in the same genomic distance were compared separately. Two sets of the discrete inputs of two biological replicates were compared using a negative binomial model, edgeR (Robinson et al., 2010). The interactions anchored at the differential H3K4me3 ChIP-seq peaks between the compared samples (p value < 0.01) were removed and only the genes with the same level of H3K4me3 peaks on promoters were processed in the downstream analysis (see Methods). **(B)** Genome browser snapshots of regions around *Hoxa* gene cluster (top) and *Sox2* (bottom). The arcs show the changes of chromatin contacts on active elements and promoters identified by the differential interaction analysis between WT ESCs and NPCs (*Hoxa* gene cluster), and the analysis between WT ESCs and MLL3/4 dCD ESCs (*Sox2*). The colors of arcs indicate degrees of interaction change between the conditions (blue to red, −/+log_10_(p-value)). The promoter regions of these genes and interacting candidate enhancer regions are shown in green and yellow shadows, respectively. H3K4me1, H3K27ac, H3K4me3, H3K27me3 ChIP-seq and RNA-seq datasets are also shown. **(C)** Genome browser snapshots of regions around *Sox2* gene. The arcs show the chromatin contacts on active elements and promoters in WT and MLL3/4 dCD cells in ESCs and NPCs. The color of arcs indicates normalized contact counts. The gene promoter and interacting candidate enhancer regions are shown in green and yellow shadows, respectively. H3K4me1, H3K27ac, H3K4me3, ChIP-seq and RNA-seq in WT and MLL3/4 dCD cells in ESCs and NPCs are also shown.

**Figure S3.**
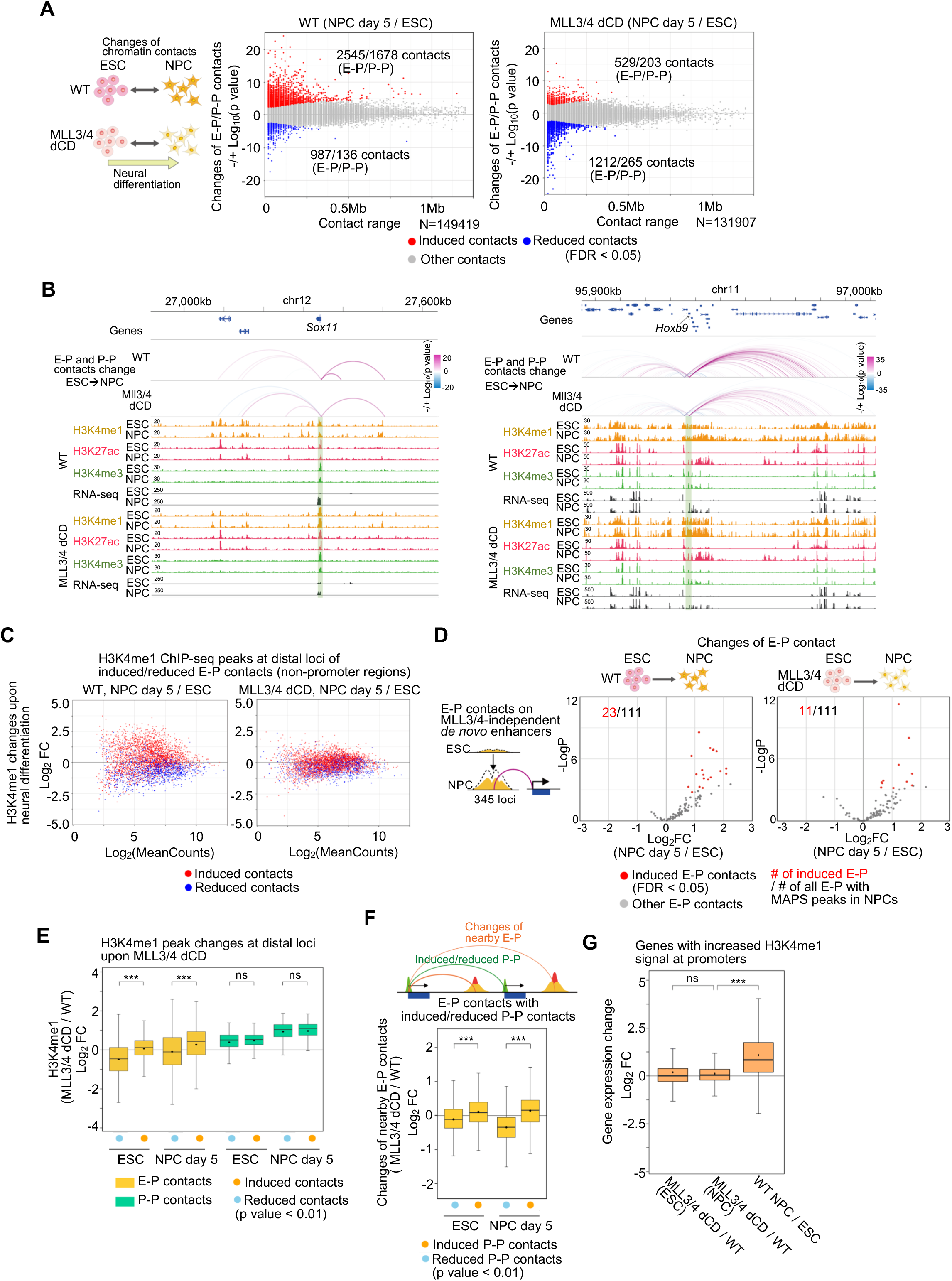
Changes of enhancer-promoter and promoter-promoter contacts upon loss of MLL3/4 catalytic activity. Related to Figure 2. **(A)** Scatter plots showing genome-wide changes of chromatin contacts anchored on promoters and enhancers (y-axis) identified in differential interaction analysis. The comparison between WT ESCs and NPCs (left) and the comparison between MLL3/4 dCD ESCs and NPCs (right) are shown. Genomic distances between their two loop anchor sites are plotted on *x*-axis. The interaction changes are indicated by significance value (−/+log_10_(p-value)). The numbers of significantly changed enhancer-promoter (E-P) and promoter-promoter (P-P) contacts are also indicated. Significantly induced and reduced chromatin contacts are shown as red and blue dots, respectively (FDR < 0.05). For details on the differential interaction analysis, see Methods and Figure S2A. **(B)** Genome browser snapshots of regions around *Sox11* gene (left) and *Hoxb9* gene (right) that failed to be activated upon NPC differentiation in MLL3/4 dCD cells. The arcs show the changes of chromatin contacts on active elements and promoters identified in differential interaction analysis between ESCs and NPCs in WT and MLL3/4 dCD cells. The colors of arcs indicate degrees of interaction change between the conditions (blue to red, −/+log_10_(p-value)). The promoter regions of these genes are shown in green shadows. H3K4me1, H3K27ac, H3K4me3 ChIP-seq and RNA-seq in ESCs and NPCs (day 5) in WT and MLL3/4 dCD cells are also shown. **(C)** Scatter plots showing changes of H3K4me1 ChIP-seq peak levels on distal element loci of significantly induced (red) and reduced (blue) E-P contacts during neural differentiation in WT cells (left). Changes of H3K4me1 ChIP-seq peak levels in MLL3/4 dCD cells on the same loci are also shown on the right. **(D)** Volcano plots showing changes of E-P contacts anchored on the MLL3/4-independent *de novo* enhancers (345 loci) upon cell differentiation from ESCs towards NPCs in WT (left) and MLL3/4 dCD cells (right). E-P contacts that were overlapped with significant peaks called by MAPS are plotted (FDR < 0.01). Significantly induced E-P contacts upon cell differentiation (FDR < 0.05) are plotted as red dots and the numbers of them are also indicated. **(E)** Boxplots of yellow boxes showing changes of H3K4me1 ChIP-seq peak signals at the distal loci of the induced and reduced E-P contacts (p value < 0.01) upon loss of MLL3/4 catalytic activity in ESCs and NPCs. Boxplots of green boxes showing changes of H3K4me1 ChIP-seq peak signals at the distal promoters of the induced and reduced P-P contacts (p value < 0.01) upon loss of MLL3/4 catalytic activity in ESCs and NPCs. ns p value > 0.05, *** p value < 0.001, two-tailed t-test. **(F)** Boxplots showing the changes of nearby E-P contacts that were anchored on the induced and reduced P-P contacts (p value < 0.01) upon loss of MLL3/4 catalytic activity in ESCs and NPCs (Schematic representation on the top). Changes of nearby E-P contacts anchored on induced and reduced P-P contacts were compared in each time point. *** p value < 0.001, two-tailed t-test. **(G)** Boxplots showing gene expression changes of genes with significantly increased H3K4me1 ChIP-seq peaks around their TSS (≤ 10 kb) upon loss of MLL3/4 catalytic activity in ESCs (left) and NPCs (middle). Gene expression changes of genes with significantly increased H3K4me1 ChIP-seq peaks around their TSS (≤ 10 kb) upon NPC differentiation in WT cells were also plotted (right). *** p value < 0.001, ns p value > 0.05, two-tailed t-test.

**Figure S4.**
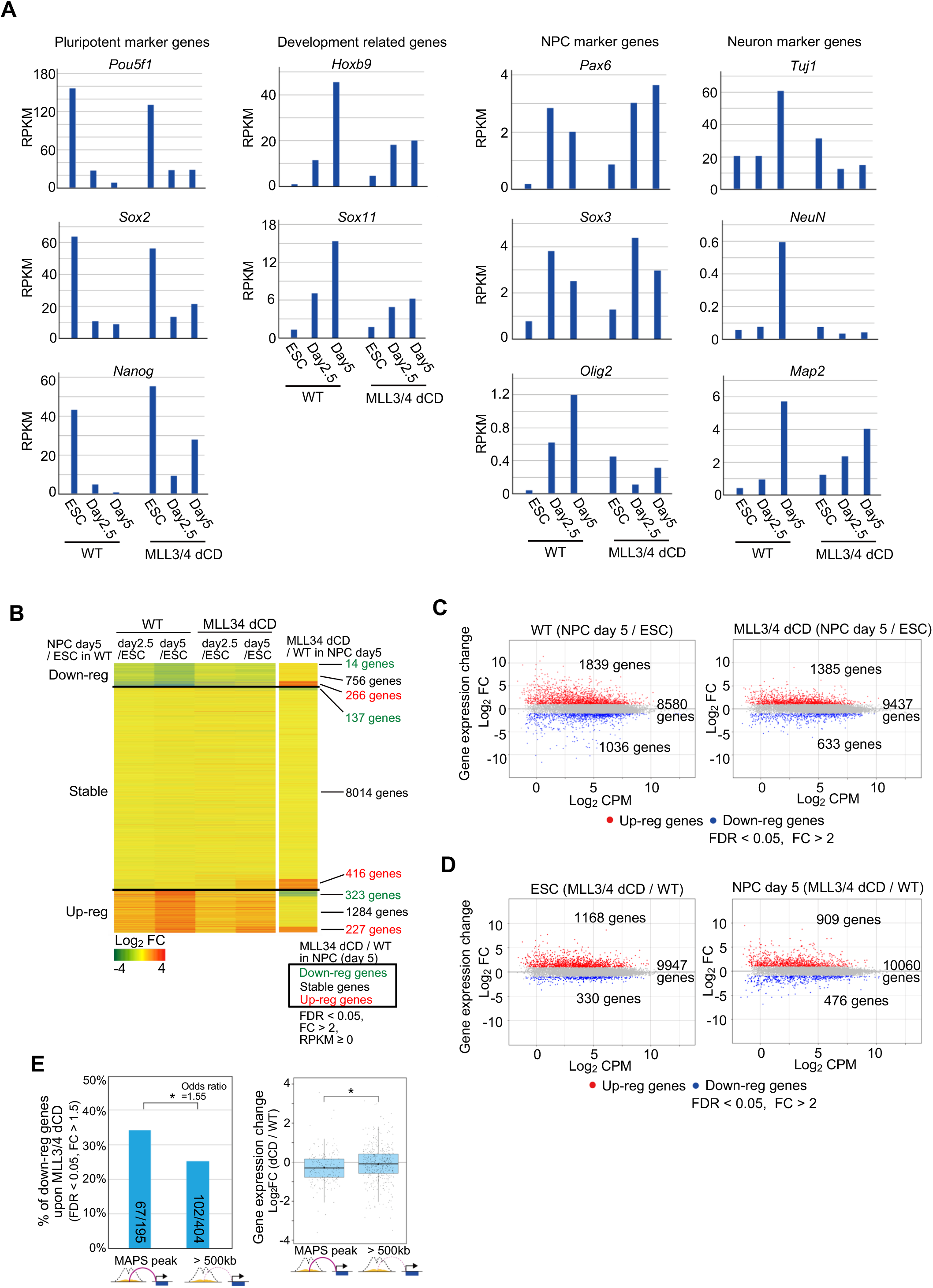
Changes of gene expression profiles upon loss of MLL3/4 catalytic activity. Related to Figure 3. **(A)** Gene expression levels of pluripotent marker genes (*Pou5f1*, *Sox2*, *Nanog*) and development related genes (*Hoxb9*, *Sox11*), and gene expression levels of NPC (*Pax6*, *Sox3*, *Olig2*) and neuron (*Tuj1*, *NeuN*, *Map2*) marker genes during neural differentiation. RPKM (reads per kilobase of transcript, per million mapped reads) values of RNA-seq (average of two replicates) at each time point in WT and MLL3/4 dCD cells are shown. **(B)** Heatmaps showing the changes of gene expression levels between ESCs and NPCs (day 2.5 and 5) in WT and MLL3/4 dCD cells (4 columns on the left), and the changes between WT and MLL3/4 dCD cells in day 5 NPCs (right column). Genes were classified based on the gene expression changes between ESCs and day 5 NPCs in WT cells (FC > 2, FDR < 0.05) and further classified based on the changes between WT and MLL3/4 dCD cells in day 5 NPCs (FC > 2, FDR < 0.05). The numbers of genes in each group (3 x 3 = 9 groups) are also shown. **(C)** Gene expression changes between ESCs and NPCs (day 5) in WT (left) and MLL3/4 dCD cells (right). Differentially up-regulated and down-regulated genes are plotted in red and blue, respectively (FC > 2, FDR < 0.05). The total number of up-regulated and down-regulated genes are also indicated. **(D)** Gene expression changes between WT and MLL3/4 dCD cells in ESCs (left) and NPCs (day 5) (right). Differentially up-regulated and down-regulated genes are plotted as red and blue dots, respectively (FC > 2, FDR < 0.05). The total number of up-regulated and down-regulated genes are indicated. **(E)** Histogram (left) shows the fraction of down-regulated genes (FC > 1.5, FDR < 0.05) in genes shown in Figure 3F (195 genes) and genes that were located at over 500-kb genomic distance away from the MLL3/4-dependent *de novo* enhancers. * p value < 0.05, Fisher’s exact test. Boxplots (right) shows fold changes of the gene expression levels of these group genes. Central bar, median; lower and upper box limits, 25th and 75th percentiles, respectively; whiskers, minimum and maximum value within the range of (1st quartile-1.5*(3rd quartile- 1st quartile)) to (3rd quartile+1.5*(3rd quartile- 1st quartile)). * p value < 0.05, two-tailed t-test.

**Figure S5.**
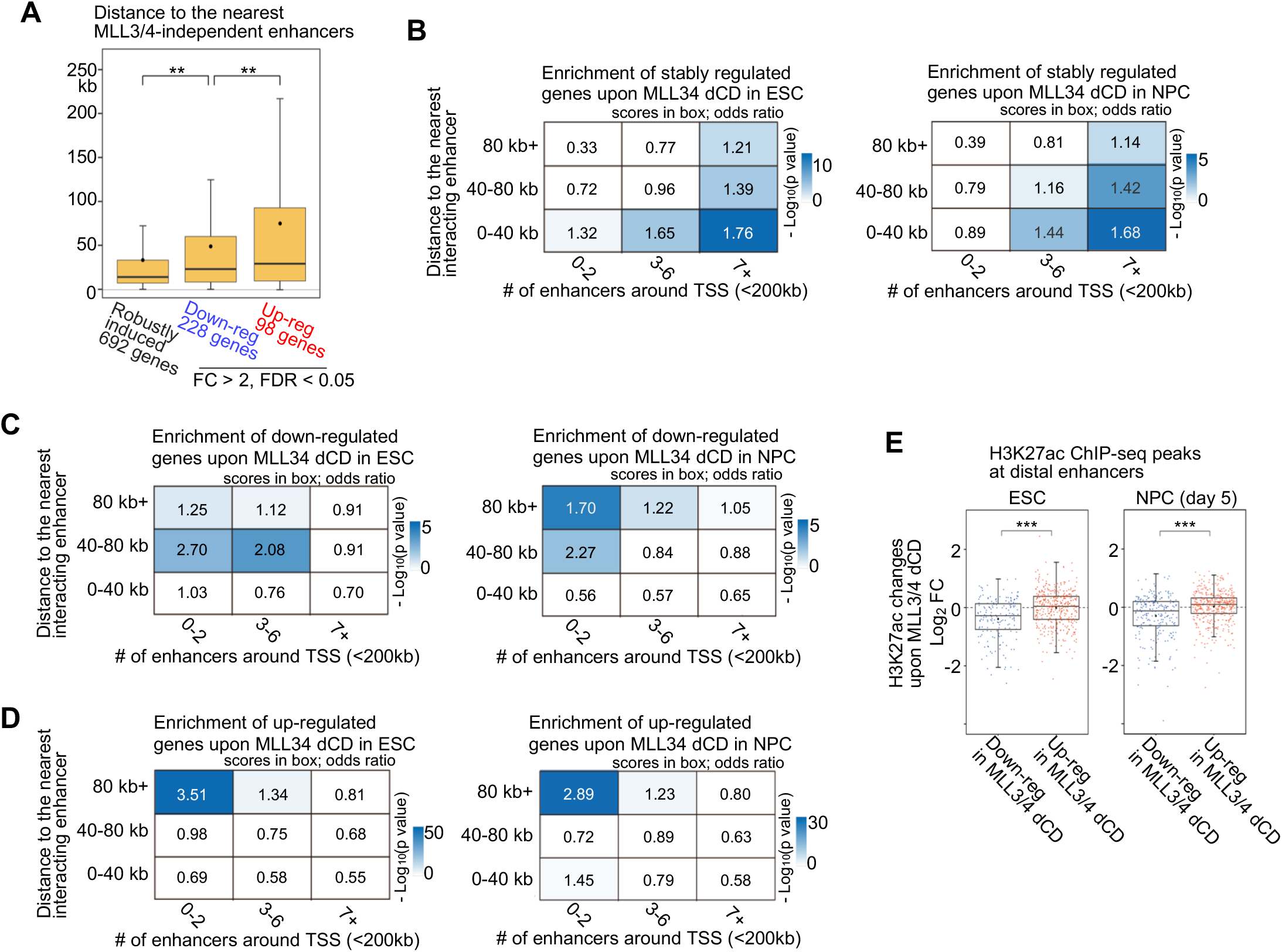
Features of MLL3/4 catalytic activity-dependent and -independent genes. Related to Figure 4. **(A)** Boxplots showing the genomic distance to the nearest MLL3/4-independent candidate enhancers from each group gene. These NPC-differentiation induced genes were classified based on the differential gene expression analysis in Figure 3C. Central bar, median; lower and upper box limits, 25th and 75th percentiles, respectively; whiskers, minimum and maximum value within the range of (1st quartile-1.5*(3rd quartile- 1st quartile)) to (3rd quartile+1.5*(3rd quartile- 1st quartile)). ** p value < 0.01, one-tailed t-test. **(B–D)** Enrichment analysis of stably expressed genes that were not differentially expressed in MLL3/4 dCD ESCs (left) and NPCs (right) (defined in Figure S4D) (B). Genes were categorized based on the distance to the nearest interacting enhancer (vertical columns) and the number of enhancers around TSS (< 200 kb) (horizontal columns). Enrichment values are shown by odds ratio (scores in boxes) and p-values (color). The distance to the nearest interacting enhancer is represented by the shortest genomic distance of significant PLAC-seq peaks on enhancers and promoters (p-value < 0.01). Enrichment analysis of the other significantly down-regulated and up-regulated genes (FC > 2, FDR < 0.05) in MLL3/4 dCD ESCs and NPCs are also shown in panel C and D, respectively. For details on the odds ratio calculation and statistical analysis, see Methods. **(E)** Boxplots showing changes of H3K27ac ChIP-seq peak signals (dCD/WT) at the all interacting distal enhancers (MAPS, p value < 0.01) of down-regulated genes and up-regulated genes in MLL3/4 dCD ESCs (left) and NPCs (right) (defined in Figure S4D). *** p value < 0.001, two-tailed t-test.

## SUPPLEMENTARY TABLES

**Table S1.**

List of NGS sample information. Related to Figures 1–4.

**Table S2.**

List of MLL3/4 catalytic activity-dependent and -independent candidate enhancers in NPCs. Related to Figure 1.

**Table S3.**

List of differentially changed enhancer-promoter contacts. Related to Figure 2.

**Table S4.**

Gene expression changes during neural differentiation in WT and MLL3/4 dCD cells. Related to Figures 3 and 4.

